# CXCR6 marks polyfunctional effector CD4 T cells required for anti-*Chlamydia* immunity in the female reproductive tract

**DOI:** 10.64898/2026.07.05.736571

**Authors:** Miguel A.B. Mercado, Yejin Kim, Qiang Li, Lin-Xi Li

**Author notes:** Correspondence should be addressed to: Miguel A.B. Mercado. 3333 Burnet Ave, Cincinnati, OH 45229.

## Abstract

CD4 T cells are essential for protective immunity against *Chlamydia* in the female reproductive tract (FRT), yet the characteristics of protective mucosal effector CD4 T cells remain poorly defined. We previously identified the transcription factor BHLHE40 as a key regulator of polyfunctional effector CD4 T cell differentiation during *Chlamydia* infection. Here, we identify the chemokine receptor CXCR6 as a marker of these protective T cells. Following intravaginal *Chlamydia muridarum* infection, *Bhlhe40*-deficient mice exhibited reduced frequencies of CXCR6⁺ CD4 T cells that correlated with impaired bacterial control. CXCR6 expression on T cells was associated with loss of stem-like features and acquisition of an effector phenotype. Compared with CXCR6⁻ cells, CXCR6⁺ CD4 T cells displayed enhanced proliferation and polyfunctionality by co-producing cytokines IFN-γ, IL-17A, and GM-CSF. Although CXCR6 was dispensable for CD4 T cell homing to the FRT, it promoted localization to the infected epithelium and the emerging memory lymphoid clusters. Importantly, depletion of CXCR6⁺ CD4 T cells reduced polyfunctional effectors and impaired bacterial clearance. Collectively, these findings identify CXCR6 as a marker of protective polyfunctional CD4 T cells and implicate CXCR6-dependent tissue positioning as a key component of effective mucosal immunity, highlighting CXCR6 as a potential biomarker for Chlamydia vaccine development.

## Introduction

Protective adaptive immunity against the obligate intracellular bacterium *Chlamydia* in the female reproductive tract (FRT) depends on CD4 T cells, yet the features that distinguish protective from non-protective CD4 T cells remain poorly defined. Early work centered on IFN-γ-producing Th1 cells as the principal mediators of anti-*Chlamydia* immunity in both mice and humans. However, recent studies demonstrated that neither IFN-γ nor the Th1 master transcription factor T-bet appear to be required for reducing *Chlamydia muridarum* burden in the mouse FRT ^1–3^. Similarly, Th17 responses have been reported to exert either protective or pathogenic effects ^4–6^. Increasing evidence suggest that anti-*Chlamydia* immunity cannot be fully explained by a single T helper lineage or signature cytokine, highlighting a need to define the cellular and functional characteristics of CD4 T cells responsible for antibacterial activity ^7,8^.

We recently reported that mice lacking the transcription factor BHLHE40 exhibit increased bacterial burden and delayed clearance of *Chlamydia muridarum* from the FRT ^9^. Impaired protection in *Bhlhe40*-deficient mice was associated with an expansion of TCF1^+^ stem-like CD4 T cells and IL-10-producing type 1 regulatory T (Tr1) cells, accompanied by a reduction in polyfunctional effector CD4 T cells capable of co-producing multiple cytokines, including IFN-γ, IL-17A and GM-CSF. These findings suggest that protective immunity is determined not only by T helper lineage specification but also by the differentiation state and functional capacity of pathogen-specific CD4 T cells. Defining markers that identify these protective polyfunctional effectors may therefore provide mechanistic insights into anti-*Chlamydia* immunity and inform vaccine development. Such studies are particularly relevant given the urgent need of a licensed human Chlamydia vaccine, as *Chlamydia trachomatis* remains to be the most frequently reported bacterial sexually transmitted infection in the United States and a major cause of pelvic inflammatory disease, ectopic pregnancy, and infertility ^10–12^.

Chemokine receptors are dynamically regulated during T cell activation, differentiation and tissue migration, and can delineate functionally distinct immune populations within mucosal tissues ^13^. During *Chlamydia* infection, several chemokine receptors, including CXCR3 and CCR5, together with the integrin α4β1, contribute to CD4 T cell trafficking and protective immunity in the FRT ^14,15^. In contrast, CCR7 restrains protective responses, as CCR7-deficient mice exhibit accelerated bacterial clearance ^16^. More recently, we identified CX3CR1 as a marker of a small subset of activated CD4 T cells in the FRT with cytotoxic characteristics ^9^. Whether protective polyfunctional CD4 T cells are similarly associated with a distinct chemokine receptor profile remains unknown.

In this study, we investigated CXCR6 (CD186/BONZO/STRL33), the sole receptor for CXCL16 ^17^, as a candidate marker of protective polyfunctional CD4 T cells during *Chlamydia* infection. CXCR6 is expressed by activated Th1, Th2, and Th17 cells and is widely used as a marker of tissue-resident memory CD8 T cells ^18–22^, yet its role in CD4 T cell-mediated immunity in the FRT remains unclear. Using the mouse model of *Chlamydia muridarum* intravaginal infection, we found that *Bhlhe40*-deficient mice exhibited reduced frequencies of CXCR6⁺ CD4 T cells, correlating with impaired bacterial control. CXCR6 marked highly proliferative effector CD4 T cells with enhanced polyfunctional cytokine production. Although CXCR6 was dispensable for CD4 T cell homing to the FRT, CXCR6⁺ CD4 T cells preferentially localized to the infected epithelium and developing memory lymphoid clusters, suggesting a role in positioning effector cells within key mucosal niches. Moreover, antibody-mediated depletion of CXCR6⁺ CD4 T cells increased bacterial burden and delayed clearance. Together, our findings establish CXCR6 as a marker of protective polyfunctional CD4 T cells and provide insight into how effector differentiation and tissue organization contribute to mucosal immunity against *Chlamydia*.

## Materials and Methods

### Mice

B6 (C57BL/6, JAX #000664), CD45.1 (B6.SJL-*Ptprc^a^Pepc^b^*/BoyJ, JAX #002014), and *Bhlhe40*^-/-^ (B6.129S1(Cg)-*Bhlhe40^tm1.1Rhli^*/MpmJ, JAX #029732) mice were purchased from The Jackson Laboratory. *Chlamydia*-specific TCR-transgenic (TP1) mice were provided by Drs. Taylor Poston and Toni Darville (UNC Chapel Hill) via Dr. Stephen McSorley (UC Davis) ^23^. All mice used for experiments were 6 to 24 weeks old. Mice were maintained under SPF conditions, and all mouse experiments were approved by the University of Arkansas for Medical Sciences Institutional Animal Care and Use Committee (IACUC).

### Chlamydia strain, mouse infection, and bacteria enumeration

*Chlamydia muridarum* strain Nigg II was originally purchased from ATCC (VR-123; Manassas, VA). *C. muridarum* was propagated in McCoy cells, elementary bodies (EBs) purified by discontinuous density gradient centrifugation, and titrated on HeLa 229 cells as previously described ^24^. Mice were synchronized for estrous by subcutaneous injection of 2.5 mg medroxyprogesterone (Depo-Provera, Greenstone, NJ) 5-7 days prior to intravaginal infection. For intravaginal (i.vag.) infection, 1×10^5^ *C. muridarum* in SPG buffer was deposited directly into the vaginal vault using a pipet tip. To enumerate bacterial shedding from the lower FRT, vaginal swabs were collected, suspended in SPG buffer, and disrupted with glass beads. Inclusion forming units (IFUs) were determined by plating serial dilutions of swab samples on HeLa 229 cells, staining with anti-MOMP mAb (clone Mo33b), and counting under a microscope.

### Flow cytometry

Spleen, DLNs, and FRT were harvested, and single cell suspensions were prepared in RPMI containing 5% fetal bovine serum (FBS). FRTs were digested with collagenase IV (Sigma) at 37 °C by GentleMACS (Miltenyi Biotech), and live cells were purified using a Percoll gradient (44%/67%). For intracellular cytokine staining, cells were stimulated with PMA (50 ng/mL) and ionomycin (500 ng/mL) for 3 hrs at 37 °C before surface and intracellular staining using the BD Cytofix/Cytoperm Kit (BD Biosciences) or intranuclear staining using the Foxp3 staining kit (ThermoFisher). The following clones of anti-mouse antibodies were obtained from BioLegend: anti-CD4 (RM4-5), anti-CD8a (53-6.7), anti-CD11b (M1/70), anti-CD44 (IM7), anti-CD45.1 (A20), anti-CD45.2 (104), anti-CD45R (B220; RA3-6B2), anti-CD90.1 (OX-7), anti-CD90.2 (53-2.1), anti-CXCR6 (SA051D1), anti-F4/80 (BM8), anti-GM-CSF (MP1-22E9), anti-KLRG1 (2F1/KLRG1), and anti-Ly108 (SLAMF6; 330-AJ). The following clones of anti-mouse antibodies were obtained from ThermoFisher (eBioscience): anti-IFN-γ (XMG1.2), anti-Ki67 (SolA15), anti-TCRVβ10b (B21.5), and eFluor 506 and eFluor 780 Fixable Viability Dye (FVD). The anti-IL-17A (TC11-18H10) was obtained from BD Biosciences. Flow cytometry data were collected on an LSR Fortessa (BD Biosciences), an LSR Celesta (BD Biosciences), or a Northern Lights cytometer (Cytek Biosciences). Data were then analyzed using FlowJo software (BD Biosciences). Polyfunctionality index was calculated using a previously published method ^25^.

### T cell adoptive transfer and CRISPR/Cas9 gene editing

CD4 T cells were isolated from the spleens of donor mice and purified using the MojoSort Mouse CD4 T Cell Isolation Kit (BioLegend). The purity of donor CD4 T cells were assessed using flow cytometry. For TP1 adoptive transfer, 1×10^5^ WT TP1 cells (CD90.1^+^) were isolated and transferred intravenously into B6 recipient mice. CRISPR/Cas9-mediated gene editing was carried out on purified TP1 cells using a P3 Primary Cell 4D-Nucleofector X Kit S (Lonza) as described by Nüssing and colleagues ^26^. Briefly, sgRNAs targeting murine *Cd90.1* (5’-CCUUGGUGUUAUUCUCAUGG-3’) and *Cxcr6* (5’-CAAGAGUCAGCUCUGUACGA-3’) (Synthego) were mixed with Alt-R *S.p*. Cas9 Nuclease V3 (IDT) and incubated at room temperature for 10 minutes. Purified TP1 cells were resuspended in P3 buffer, mixed with the sgRNA/Cas9 RNP complex, and electroporated on a Nucleofector (Lonza). Cells were then rested for 10 minutes at 37°C in a CO_2_ incubator before transferred intravenously into recipient mice. All recipient mice were challenged intravaginally with *C. muridarum* one day after adoptive transfer.

### Immunofluorescence imaging

FRTs were isolated and fixed with 4% PFA for 2 hrs at 4°C, washed with cold PBS, and incubated in 30% sucrose overnight at 4°C. Whole tissues were then embedded in O.C.T. and 7 µm frozen sections cut using the HM 505 E Cryostat (Microm). Slides with tissue sections were fixed with acetone at -20°C for 15 minutes. Before staining, frozen sections were rehydrated in PBS, blocked with 5% BSA, and treated with Streptavidin/Biotin Blocking Kit (VectorLabs) for 30 minutes. Primary and secondary staining were performed for 1 hr in the dark, followed by DAPI staining for 15 minutes at room temperature. Sections were then mounted with Fluoromount-G Mounting Medium (SouthernBiotech). All images were captured and processed with a BZ-X800 Microscope (Keyence). The following antibodies from BioLegend were used for staining: Alexa Fluor 488 anti-CD4 (RM4-5), Biotin anti-CXCR6 (SA051D1), Alexa Fluor 647 Streptavidin, and 4’,6-Diamidino-2-Phenylindole, Dilactate (DAPI).

### Monoclonal Ab treatment

Ultra-LEAF Purified anti-mouse CD186 (CXCR6) depleting antibody (clone SA051D1) was custom-made by BioLegend. Antibody treatment was performed by intraperitoneal injection of 0.3 mg of anti-CXCR6 on days 6, 8, 10, 13, 15, 17, 20, 22, 24, and 27 following *C. muridarum* intravaginal infection.

### Single-cell RNA sequencing data analysis

Single-cell RNA sequencing dataset used in this study were previously published and are available through the NCBI Gene Expression Omnibus (GEO) under accession number GSE253394 ^9^. Data analysis was conducted using Seurat v.4.3 under R v4.3.1 environment.

### Statistical analysis

Statistical analysis was performed using Prism 10 (GraphPad Software). An unpaired *t* test or a paired *t* test was used for normally distributed continuous-variable comparisons. A Mann-Whitney U test was used for nonparametric comparisons. For the comparison of multiple groups, the ANOVA tests were used followed by multiple comparisons of means. A *p*-value < 0.05 was considered statistically significant.

## Results

### Reduced CXCR6+ CD4 T cells in the FRT of Bhlhe40-deficient mice following intravaginal Chlamydia infection

In a previous study, we reported that mice lacking the transcription factor BHLHE40 in CD4 T cells are more susceptible to *Chlamydia* infection in the FRT than WT controls ^9^. In our published single-cell RNA sequencing dataset, we showed that *Bhlhe40*-deficient CD4 T cells were enriched in clusters that resemble *Tcf7*^+^ stem-like CD4 T cells (#3, #10) and Tr1 cells (#6), whereas WT counterparts displayed signatures of robust TCR signaling (#0), T-helper polyfunctionality (#4), and cytotoxicity (#8) (**Fig. 1A**). These transcriptomic data prompted us to identify surface markers uniquely expressed on protective polyfunctional CD4 T cells in the FRT, which would potentially serve as biomarkers of protective immunity to inform vaccine development. Among surface molecules examined, we found that *Cxcr6* (gene encoding chemokine receptor CXCR6) was highly expressed in WT-enriched CD4 T cell clusters but not in stem-like CD4 T cells (#3, #10) enriched in *Bhlhe40*-deficient FRT or regulatory T cells (#9) (**Fig. 1B**). Flow cytometry analysis confirmed that the frequency of CXCR6+ CD4 T cells was significantly lower in *Bhlhe40*-deficient mice compared to WT controls at 14 days post *C. muridarum* intravaginal infection (**Fig. 1C and 1D**). Moreover, CXCR6 expression was accompanied by decreased SLAMF6 and increased KLRG1 expression on the cell surface, which is consistent with the loss of stemness and gain of an effector-like phenotype, respectively, of these CXCR6+ CD4 T cells in the FRT (**Fig. 1C-1F**). Together, these findings identify CXCR6 as a marker associated with differentiated effector CD4 T cells that are diminished in the absence of BHLHE40.

**Fig 1.**
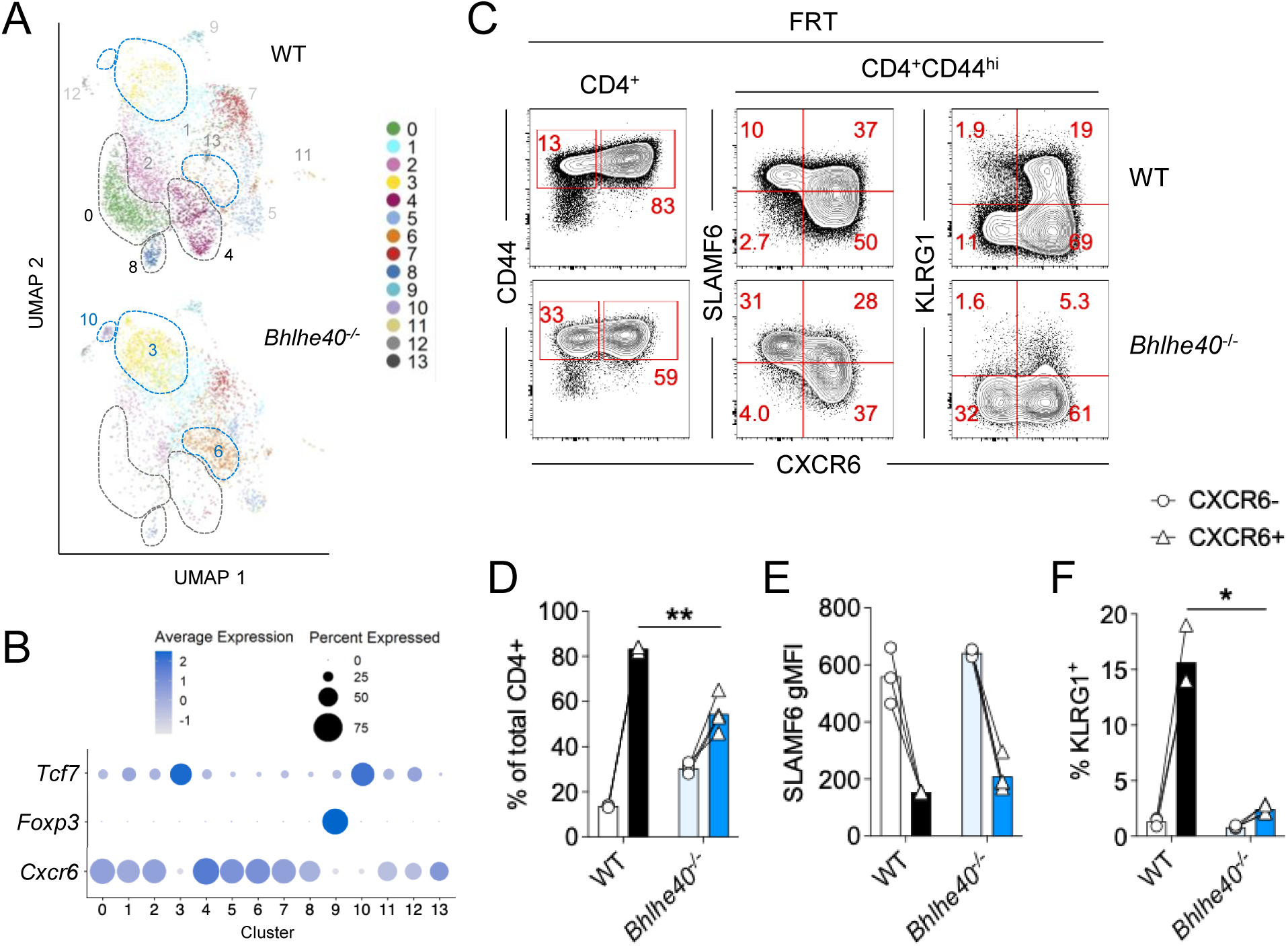
*Bhlhe40*^-/-^ mice have less CXCR6^+^ CD4 T cells in the FRT than WT mice following intravaginal *Chlamydia muridarum* infection. WT and *Bhlhe40*^-/-^ mice were infected intravaginally with 1×10^5^ *C. muridarum.* CD4 T cells in the female reproductive tract (FRT) were analyzed at 14 days post infection (dpi) by single-cell RNA sequencing (A-B) or flow cytometry (C-F). **(A)** Deconvoluted UMAP depicting 14 clusters of activated CD4 T cells (live CD90.2^+^CD4^+^CD44^hi^) from WT and *Bhlhe40^-/-^* FRT (*Mercado et al.*, 2024, *PLOS Pathogens;* reproduced under CC BY). **(B)** Dot plot depicting *Tcf7, Foxp3,* and *Cxcr6* mRNA expression in each cluster in (A). **(C)** Representative FACS plots depicting total or activated (CD44^hi^) CD4 T cells in the FRT (gated on live CD11b^-^B220^-^F4/80^-^CD90.2^+^CD8^-^ cells). **(D)** Percentages of CXCR6^-^ and CXCR6^+^ cells within total CD4 T cells in the FRT. **(E)** Geometric mean fluorescence intensities (gMFI) of SLAMF6 in CD44^hi^ CXCR6^+/-^ CD4 T cells. **(F)** Percentages of KLRG1^+^ cells in CD44^hi^ CXCR6^+/-^ CD4 T cells. Data represents two similar experiments with 3-4 mice per group. Each pair of data points represents the CXCR6^-^ and CXCR6^+^ CD4 T cells within the same mouse. \**p* < 0.05, \*\**p* < 0.01 by paired *t* tests.

### CXCR6+ CD4 T cells display enhanced effector function and increased proliferation compared to CXCR6- CD4 T cells in the FRT

Earlier studies have defined CXCR6 as a marker for highly differentiated type 1 T cells with tissue homing potential in human PBMC ^18^. In autoimmune models, CXCR6 expression on CD4 T cells is associated with *Il17a* and *Csf2* upregulation ^20^. To determine whether CXCR6+ CD4 T cells resemble the polyfunctional CD4 T cell population capable of producing multiple cytokines in the context of *Chlamydia* infection, we measured IFN-γ, IL-17A and GM-CSF production by CD4 T cells in the FRT 14 days following *C. muridarum* intravaginal infection. These three cytokines were identified as the major cytokines produced by polyfunctional CD4 T cells in our previous study ^9^. As shown in **Fig. 2A-C**, CXCR6+ CD4 T cells contained a higher frequency of IFN-γ+ cells and produced more IFN-γ on a per-cell basis than the CXCR6- counterparts. Moreover, CD4 T cells producing multiple cytokines (IFN-γ+IL-17A+, IFN-γ+GM-CSF+, and IFN-γ+IL-17A+GM-CSF+) were preferentially enriched within the CXCR6+ compartment, whereas CXCR6-cells were predominantly IFN-γ single producers (**Fig. 2D**). These differences were reflected by a significantly higher polyfunctionality index among CXCR6+ CD4 T cells (**Fig. 2E**). Notably, within the CXCR6+ population, we found that polyfunctional CD4 T cells co-producing 2 or 3 cytokines expressed higher levels of surface CXCR6 on a per-cell basis than IFN-γ single producers (**Fig. 2F**), indicating a possible trajectory of acquiring polyfunctionality while upregulating CXCR6 expression during effector CD4 T cell maturation. Finally, CXCR6+ CD4 T cells in the FRT contained slightly higher % of Ki67+ cells than CXCR6- counterpart (**Fig. 2G-H**), indicating that increased proliferation may contribute to the enrichment of CXCR6+ cells in the FRT. These results demonstrate that CXCR6 expression identifies a highly proliferative population of polyfunctional effector CD4 T cells in the infected FRT.

**Fig 2.**
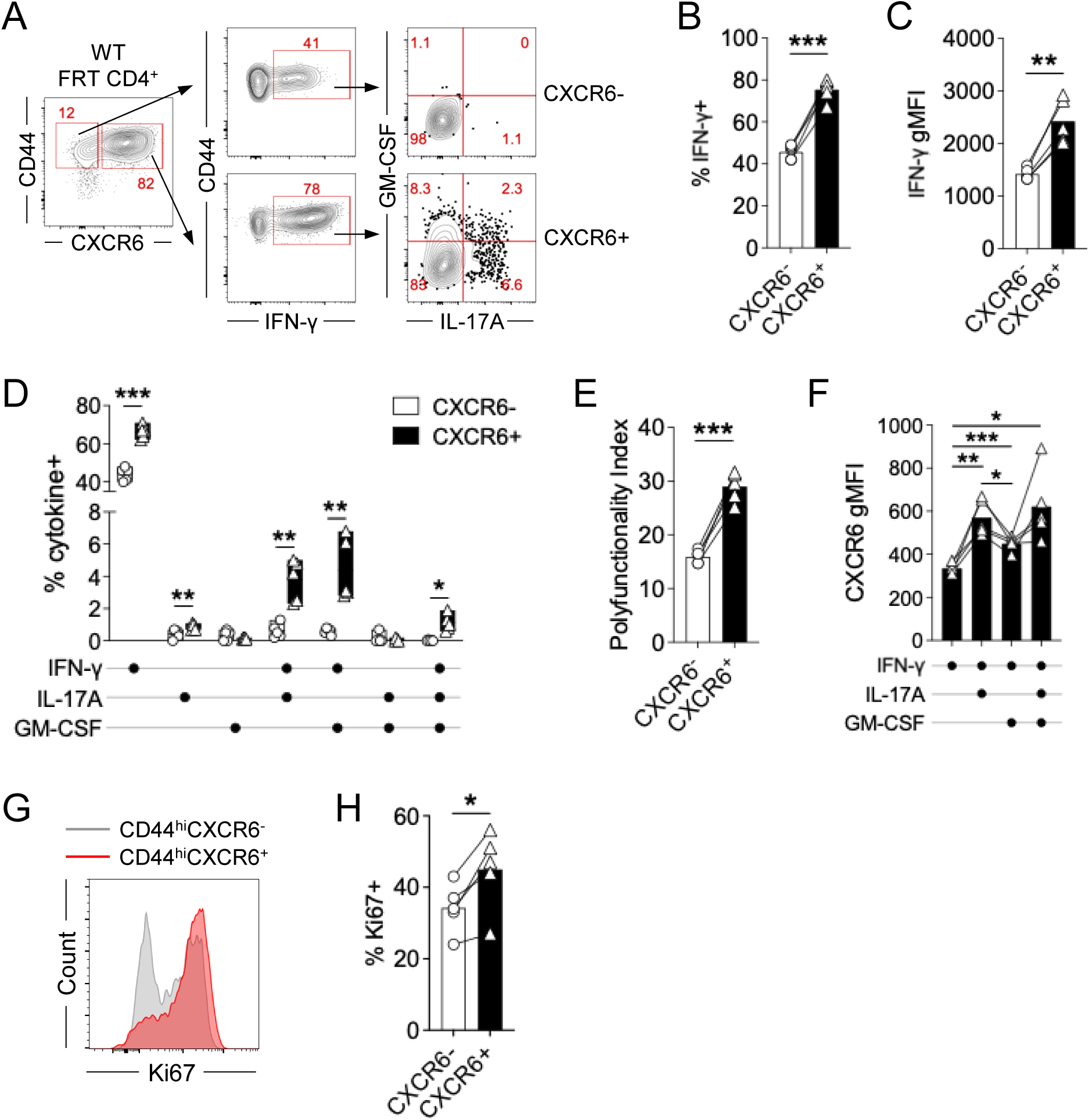
CXCR6^+^ CD4 T cells display enhanced cytokine production and augmented proliferation compared to CXCR6^-^ CD4 T cells in the FRT. B6 mice were infected intravaginally with 1×10^5^ *C. muridarum.* CD4 T cells in the FRT were analyzed at 14 dpi. **(A)** Representative FACS plots depicting IFN-γ-, IL-17- and GM-CSF-producing CD4 T cells within the CD44^hi^ CXCR6+/- populations in the FRT (gated on live CD90.2^+^CD4^+^ cells). **(B-D)** Summary data depicting percentages of IFN-γ+ cells (B), geometric mean fluorescence intensity (gMFI) of IFN-γ (C), percentages of IFN-γ-, IL-17A-, and GM-CSF-producing polyfunctional T cells (D) in CD44^hi^ CXCR6+/- CD4 T cells. **(E)** Polyfunctionality index of CD44^hi^ CXCR6+/- CD4 T cells. **(F)** CXCR6 gMFI of IFN-γ-producing CXCR6+ CD4 T cells. **(G-H)** Representative FACS histogram of Ki67 expression (G) and summary data depicting %Ki67+ (H) in CD44^hi^ CXCR6+/- CD4 T cells. Data represents 2-3 similar experiments with 3-5 mice per group. Each pair of data points in (B-H) represents the CXCR6- and CXCR6+ CD4 T cells within the same mouse. \**p* < 0.05, \*\**p* < 0.01, \*\*\**p* < 0.001 by paired *t* tests.

### CXCR6 is preferentially induced on antigen-specific CD4 T cells during Chlamydia infection

*Chlamydia* infection in the FRT induces robust CD4 T cell clonal expansion in the secondary lymphoid organs, followed by the migration of both antigen-specific and bystander CD4 T cells to the FRT ^24,27^. To investigate the CXCR6 expression pattern in *Chlamydia*-specific CD4 T cells, we performed adoptive transfer of *Chlamydia*-specific TCR transgenic CD4 T cells (TP1) into congenic WT hosts and characterized CD4 T cell response at the peak of clonal expansion following *C. muridarum* infection (**Fig. 3A**) ^9,23^. At day 10 post infection (10 dpi), a small proportion of activated CD44^hi^ CD4 T cells upregulated CXCR6 in the draining iliac lymph nodes (DLNs), with ∼20% TP1 cells being CXCR6+ compared to ∼5% of endogenous CD4 T cells (**Fig. 3B-C**). In contrast, vast majority of both TP1 cells and endogenous CD4 T cells express CXCR6 in the FRT, and the cytokine profiles were similar between these two populations, with more polyfunctional CD4 T cells found in the CXCR6+ populations than the CXCR6- counterparts (**Fig. 3B-D**). Notably, IFN-γ single producers were significantly more abundant in the CXCR6-compartment than the CXCR6+ counterparts in TP1 cells, which was different from the endogenous T cells. Consistent with our previous report, the most abundant polyfunctional CD4 T cells in the FRT were the cells that co-produce IFN-γ and GM-CSF (**Fig. 3D**). Finally, these cytokine profiles lead to higher polyfunctionality indices in TP1 cells compared to endogenous T cells (**Fig. 3E**). Together, these data suggest that CXCR6 expression is induced upon T cell activation in the DLNs and upregulated in polyfunctional CD4 T cells in the FRT in both antigen-specific and endogenous polyclonal T cell populations.

**Fig 3.**
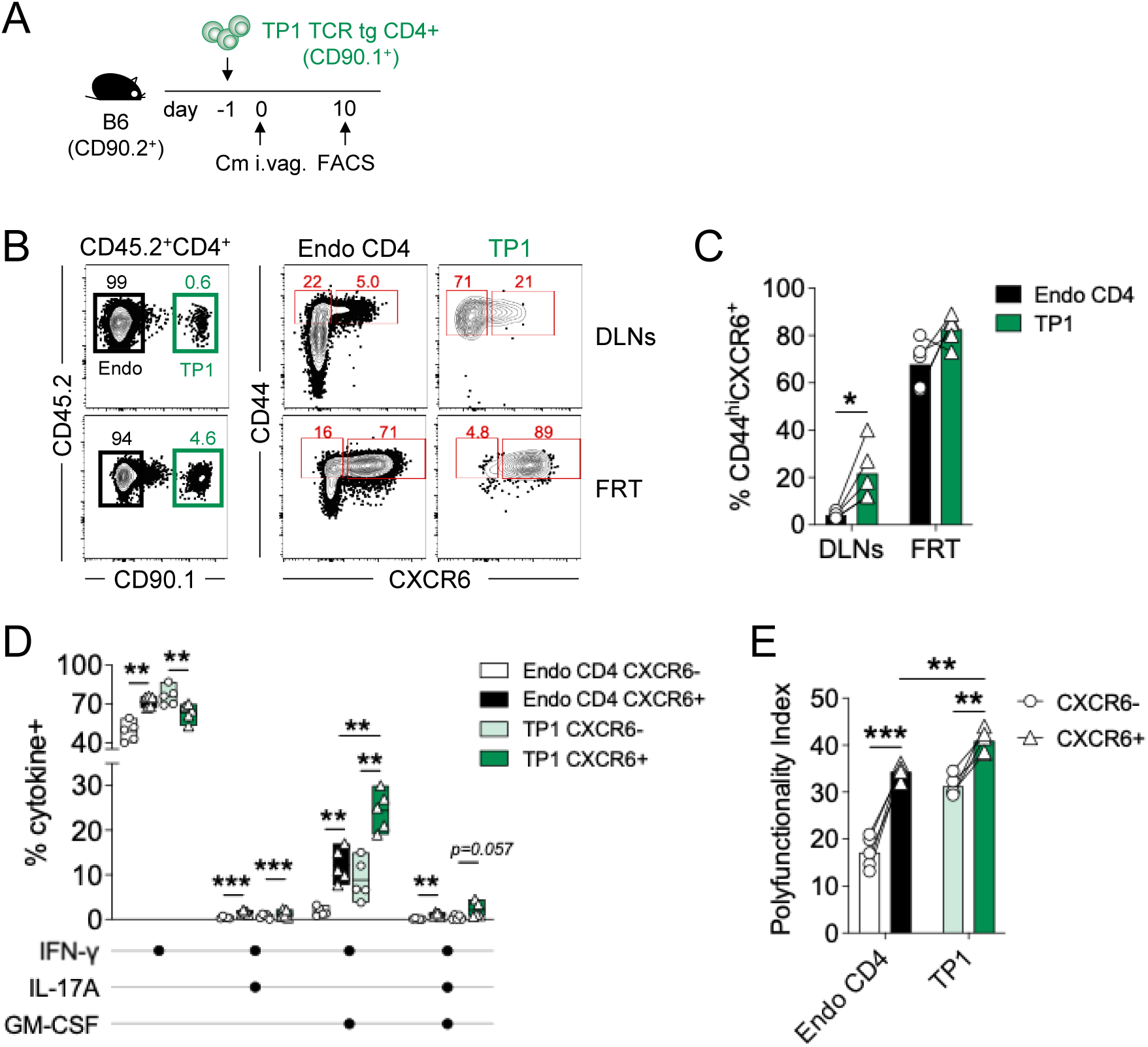
*Chlamydia*-specific CD4 T cells express higher CXCR6 and exhibit greater polyfunctionality than endogenous T cells. *Chlamydia*-specific TP1 TCR transgenic CD4 T cells (1×10^5^) were adoptively transferred into B6 hosts one day before intravaginal infection with 1×10^5^ *C. muridarum*. CD4 T cells in the FRT were analyzed at 10 dpi. **(A)** Experimental workflow. **(B-C)** Representative FACS plots (B) and summary data (C) depicting % of CD44^hi^CXCR6^+^ CD4 T cells within endogenous and TP1 CD4 T cells in the DLNs and FRT (gated on live CD45.2^+^CD4^+^ cells). **(D-E)** Summary data depicting percentages of IFN-γ-, IL-17A-, and GM-CSF-producing polyfunctional T cells (D) and polyfunctionality index (E) of endogenous and TP1 CD44^hi^ CXCR6+/- CD4 T cells in the FRT. Data represents three similar experiments with 4-5 recipient mice per experiment. Each pair of data points represents endogenous and transferred TP1 CD4 T cells (C) or CXCR6- and CXCR6+ CD4 T cells (E) within the same mouse. \**p* < 0.05, \*\**p* < 0.01, \*\*\**p* < 0.001 by paired *t* tests.

### CXCR6 is not required for CD4 T cell homing to the FRT during Chlamydia infection

Several chemokine receptors and integrins facilitate CD4 T cell homing to the FRT during *Chlamydia* infection ^14,15^. Given that CXCR6+ CD4 T cells were significantly more enriched in the FRT than the DLNs at 10 dpi (**Fig. 3C**), it is possible that CXCR6 functions as an essential chemokine receptor that facilitates CD4 T cell homing to the FRT. To test this hypothesis, we performed a competitive FRT homing assay using TP1 cells undergoing CRISPR-mediated CXCR6 deletion ^26^. TP1 cells targeted for congenic marker *Cd90.1* deletion were used as controls (**Fig. 4A**). As expected, most of the non-edited WT TP1 cells in the FRT were CXCR6+ (**Fig. 4B**). In the setting of the TP1 1:1 competitive homing, we recovered comparable numbers of CXCR6-deficient and control TP1 cells from the recipient FRT (**Fig. 4B-C**), indicating that CXCR6 is dispensable for antigen-specific CD4 T cell recruitment to the FRT following *C. muridarum* infection.

**Fig 4.**
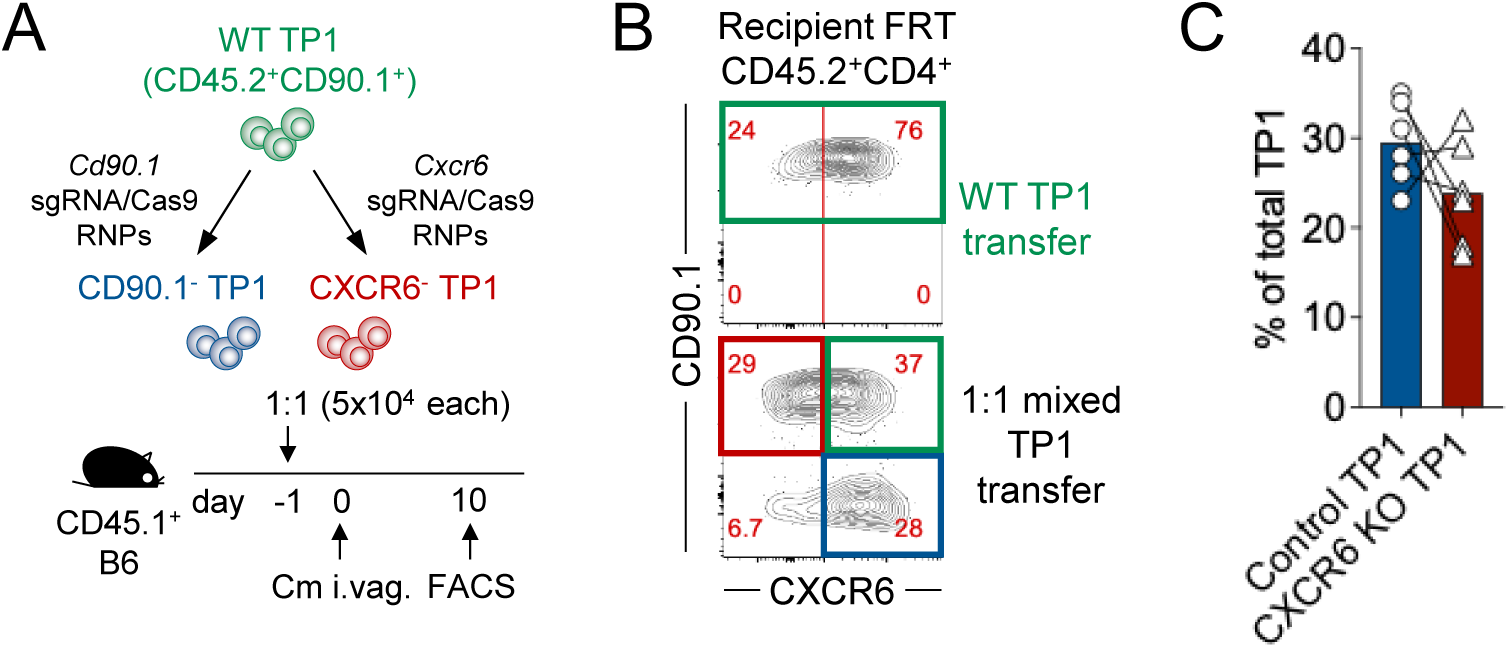
CXCR6 expression is not essential for CD4 T cell homing to the FRT during *C. muridarum* infection. Naïve *Chlamydia*-specific TP1 CD4 T cells were purified, electroporated with sgRNA/Cas9 RNPs targeting either control (*Cd90.1*) or *Cxcr6* and adoptively transferred into congenic CD45.1^+^ B6 recipient mice one day before intravaginal infection with 1×10^5^ *C. muridarum.* CD4 T cells in the FRT were analyzed at 10 dpi. **(A)** Experimental workflow. **(B)** Representative plots depicting CD90.1- and CXCR6-expressing transferred TP1 CD4 T cells in the FRT (gated on live CD45.2^+^CD4^+^ cells). Green box: WT or untargeted TP1; red box: *Cxcr6* KO TP1; blue box: control (*Cd90.1* KO) TP1. **(C)** Summary data depicting percentages of control and CXCR6 KO TP1 T cells. Each pair of data points represents control and CXCR6 KO TP1 cells within the same mouse. Data represents two similar experiments with 5-6 recipient mice per experiment.

### CXCR6 instructs CD4 T cell positioning in the FRT following Chlamydia infection

We next sought to determine whether CXCR6 was needed for CD4 T cell localization within specific regions in the FRT following tissue homing. Given that CXCL16, the ligand for CXCR6, is produced by endometrial cells and oviduct epithelial cells within the FRT ^28,29^, we hypothesized that CXCR6 may position effector CD4 T cells adjacent to the *Chlamydia*-infected epithelium. At 14 dpi, we observed a more concentrated CXCR6 immunofluorescent staining at the epithelium layer than CD4 staining, and small clusters of CXCR6-expressing CD4 T cells were positioned near epithelial cells in *Chlamydia*-infected FRT (**Fig. 5A-E**). Some CXCR6+ cells did not co-express CD4, suggesting that CXCR6-guided epithelial localization is extended to non-CD4 T cells, presumably NK cells and CD8 T cells ^21,30^. At 21 dpi, a time point when vast majority of *Chlamydia* were cleared from the FRT, and CD4 memory lymphoid clusters (MLCs) start to form ^7,31^, we observed high levels of CXCR6 expression at these CD4 MLCs (**Fig. 5F-J**). These data suggest that although CXCR6 is not required for initial homing to the FRT, it may contribute to the spatial organization of effector CD4 T cells within the infected mucosa, presumably potentiating their effector functions and memory formation at various stages of *Chlamydia* infection.

**Fig 5.**
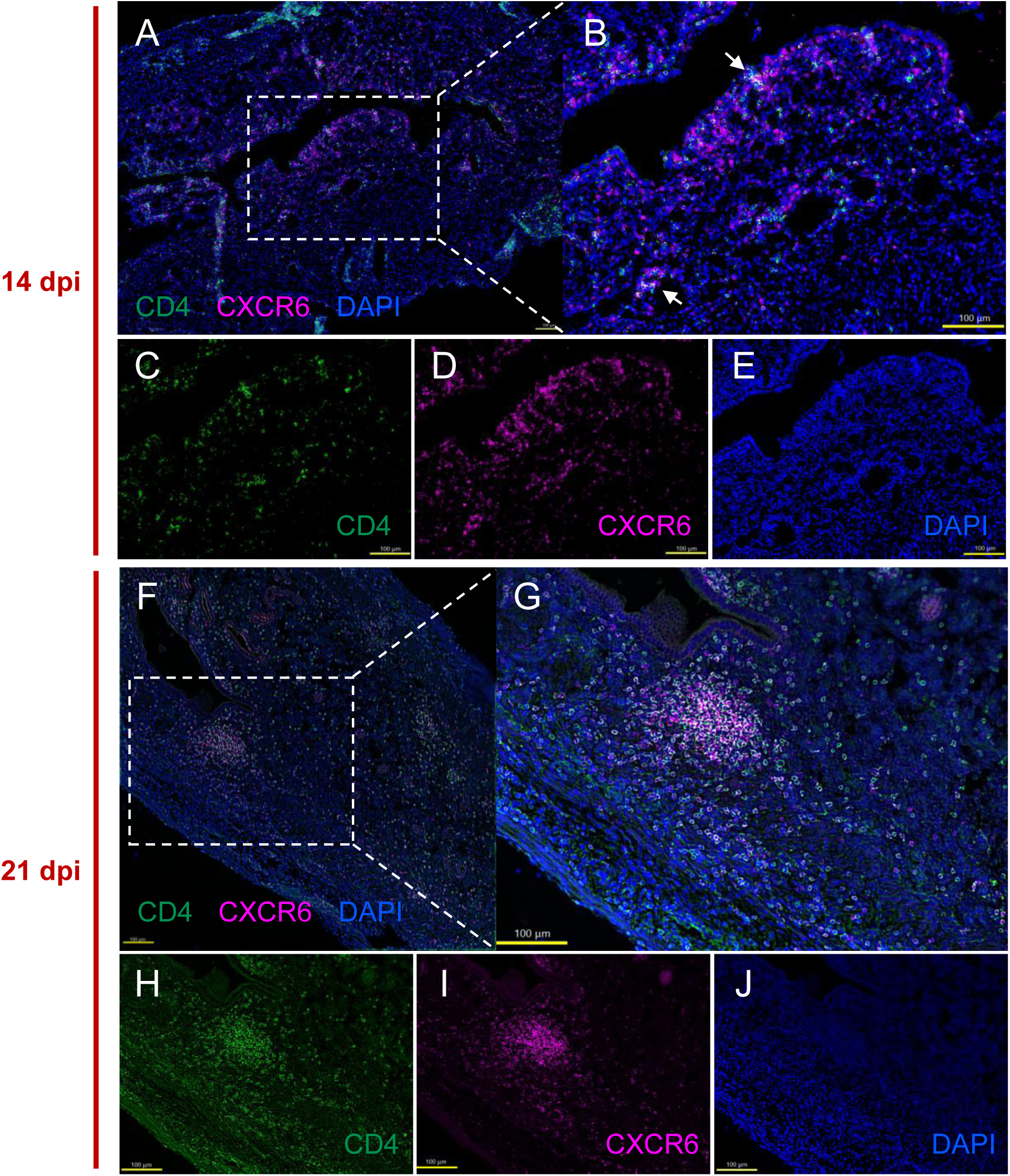
Localization of CXCR6^+^ CD4 T cells following *Chlamydia* infection. B6 mice were infected intravaginally with 1×10^5^ *C. muridarum.* FRTs were harvested at 14 (**A-E**) and 21 (**F-J**) dpi for imaging. Immunofluorescent staining of CXCR6, CD4, and DAPI in the FRT (A, F). Representative section depicting merged (B, G) and individual staining for CD4 (C, H), CXCR6 (D, I), and DAPI (E, J). Yellow scale bars, 100 μm.

### Anti-CXCR6 treatment reduces polyfunctional CD4 T cells and impairs bacterial control in the FRT

Our data so far suggest that CXCR6 marks effector CD4 T cells that are polyfunctional and uniquely positioned in the FRT following *Chlamydia* infection. Whether these CXCR6-expressing CD4 T cells are essential for *Chlamydia* control remains to be investigated. The contribution of CXCR6 to host protection varies among infectious models. CXCR6 enhances immunity during *M. tuberculosis* lung infection and anti-parasitic responses in the liver ^32,33^, whereas CXCR6-deficient mice mount normal T cell responses and control *L. monocytogenes* infection comparable to WT animals ^34^. Given that CXCR6+ CD4 T cells were significantly reduced in the *Bhlhe40*-deficient mice and protective immunity is impaired in these mice (Fig. 1 and ^9^), it is likely that CXCR6+ CD4 T cells are required for *Chlamydia* control in the FRT. To test this directly, we treated WT mice with or without anti-CXCR6 depleting antibody between days 6-27 post *Chlamydia* infection and monitored bacterial burden in the FRT (**Fig. 6A**). We chose this timeframe of treatment to minimize the impact on innate immune responses while spanning the kinetics of the CD4 T cell response. Flow cytometry analysis by the end of the treatment (29 dpi) revealed that polyfunctional CD4 T cells producing IFN-γ, IL-17A and GM-CSF were reduced in both DLNs and FRT of anti-CXCR6 treated mice (**Fig. 6B-D**). In contrast, IFN-γ single producers remained comparable between anti-CXCR6 and PBS treated groups. The selective loss of multifunctional cytokine-producing CD4 T cells resulted in diminished polyfunctionality indices in both the DLNs and FRT following anti-CXCR6 treatment (**Fig. 6E**). Importantly, we observed that anti-CXCR6 treatment resulted in heightened *Chlamydia* burdens in the FRT compared to PBS-treated controls between days 17 and 24 post infection, a critical time frame for bacterial control and clearance (**Fig. 6F**). These data suggest that CXCR6-expressing polyfunctional CD4 T cells contribute substantially to optimal bacterial control in the FRT following *Chlamydia* infection.

**Figure 6.**
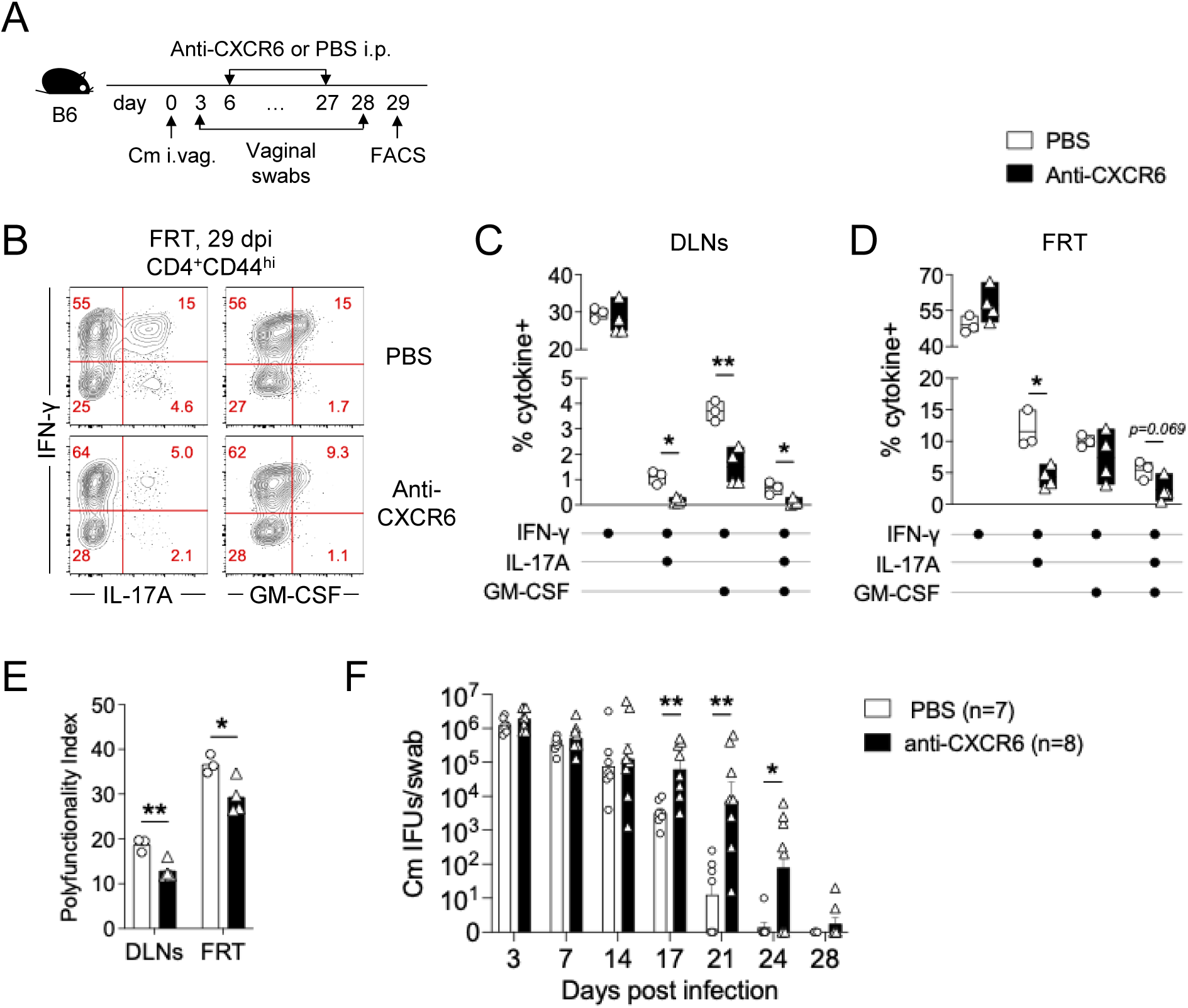
Anti-CXCR6 treatment reduces polyfunctional CD4 T cells and impairs bacterial control in the FRT. B6 mice were infected intravaginally with 1×10^5^ *C. muridarum*. Cohorts of mice were treated with anti-CXCR6 antibody or PBS between 6-27 dpi. Bacterial shedding from the lower FRT were monitored by vaginal swabs. CD4 T cells in the FRT were analyzed at 29 dpi. **(A)** Experimental workflow. **(B-E)** Representative FACS plots (B) and summary data depicting IFN-γ-, IL-17A-, and GM-CSF-producing CD4 T cells (gated on live CD90.2^+^CD4^+^CD44^hi^ cells) in the DLNs (C) and FRT (D), and polyfunctionality index (E) of anti-CXCR6- and PBS-treated mice. **(F)** Bacterial shedding from the lower FRT in anti-CXCR6- and PBS-treated mice. Data are from two combined experiments with 7-8 mice per group. Each data point in (C-E) represents an individual sample pooled from 2-3 mice. Each data point in (F) represents an individual mouse. \**p* < 0.05, \*\**p* < 0.01 by unpaired *t* tests.

## Discussion

CD4 T cells are essential for protective immunity against *Chlamydia*, with early work emphasizing IFN-γ-producing Th1 responses. Nevertheless, more recent studies suggest that protective cellular immunity is not restricted to a single CD4 T cell subset, but instead conferred by a heterogenous CD4 T cell response ^8,35,36^. In a previous study, we identified the transcription factor BHLHE40 as a regulator of protective effector CD4 T cell differentiation and polyfunctionality during *Chlamydia* primary infection. Here, we extended the prior study and identified CXCR6 as a marker for polyfunctional CD4 T cells essential for bacterial control in the female reproductive tract. Our findings suggest that CXCR6 expression delineates a differentiation state of effector CD4 T cells beyond T helper lineage commitment and highlight its role for tissue localization in CD4 T cell-mediated immunity at the FRT mucosa.

Several observations suggest that CXCR6 expression reflects an advanced stage of effector CD4 T cell differentiation rather than commitment to a particular helper lineage. CXCR6+ cells exhibited reduced SLAMF6 expression and increased KLRG1 expression, features associated with diminished stemness and terminal effector differentiation, respectively. Moreover, CXCR6+ CD4 T cells displayed enhanced proliferative activity and preferentially co-produced multiple cytokines including IFN-γ, IL-17A, and GM-CSF. Notably, substantial numbers of IFN-γ single-producing cells remained within the CXCR6- compartment, arguing against CXCR6 serving as a general marker of Th1 cells. Instead, our findings support a model in which CXCR6 identifies a highly differentiated subset of polyfunctional type 1 effector CD4 T cells. This is in agreement with an early study identifying CXCR6 expression on a subset of highly differentiated effector/memory type 1 T cells in human peripheral blood ^18^. CXCR6+ T cells are also enriched in inflamed tissues, including arthritis synovial fluid and inflamed liver, suggesting a close association between CXCR6 expression and tissue inflammation. Nevertheless, naïve T cells can be primed directly by DCs to upregulate CXCR6 in vitro, indicating that its expression is not restricted to tissue-resident populations. Consistent with this observation, CXCR6+ TP1 cells were readily detected in the DLNs (Fig. 3), suggesting that antigen-specific CD4 T cells initiate CXCR6 expression during priming and may further upregulate CXCR6 following migration into the inflamed FRT. Given that higher surface CXCR6 protein level (MFI) is associated with increased effector functions (Fig. 2F), we speculate that CXCR6 expression may serve as a continuum reflecting the differentiation state of effector CD4 T cells – this hypothesis needs to be tested experimentally.

To date, it remains unclear whether the effector function of a single cytokine is fully responsible for anti-*Chlamydia* immunity in vivo. IL-12p40, the shared subunit of cytokines IL-12 and IL-23, represents the best-characterized molecule required for bacterial control, as both IL-12p40-deficient mice and antibody-mediated IL-12 blockade exhibit impaired *Chlamydia* clearance from the FRT ^3,37,38^. Our studies have identified three major effector cytokines produced by CD4 T cells, IFN-γ, IL-17A and GM-CSF, yet deficiency of any one of these cytokines alone has little impact on *Chlamydia* control (^4,5,37^ and data not shown), suggesting substantial redundancy among individual effector pathways revealed in loss of function studies. It is possible that the superior protective capacity of CXCR6+ CD4 T cells derives from their ability to coordinately deploy multiple effector mechanisms from cytokines. Alternatively, CXCR6 may identify CD4 T cell subpopulations with effector functions other than cytokine production. In several cancer models, CXCR6 marks CD4 and CD8 T cells with cytotoxic function ^39,40^. Although not tested directly in this current study, our scRNAseq data revealed that *Cxcr6* is also highly expressed in the predicted cytotoxic CD4 T cell cluster (#8) in WT mice (Fig 1). Moreover, a recent study by Starnbach and colleagues has detected granzyme-B producing cytotoxic CD4 T cells in the FRT during *Chlamydia trachomatis* infection in mice ^41^. Together, our findings raise the possibility that CXCR6+ CD4 T cells comprise a heterogeneous population capable of mediating protection through both polyfunctional cytokine production and direct cytolytic activity against infected epithelial cells.

The contribution of CXCR6 for tissue homing and positioning appears to be context dependent. During influenza infection, CXCR6 is not required for CD8 T cell entry into the lung but is needed to position resident memory CD8 T cells in the airways as first responders to reinfection ^21^. Similarly, within the tumor microenvironment, CXCR6 acts as a key receptor that positions effector-like cytotoxic CD8 T cells near dendritic cells producing CXCL16 to receive pro-survival signals ^40^. In other contexts, however, CXCR6 is dispensable or even restrains tissue accumulation ^32,42^. Consistent with a role in tissue positioning rather than initial recruitment, CXCR6 was dispensable for CD4 T cell homing to the FRT in our model. Nevertheless, CXCR6+ CD4 T cells preferentially localized near the epithelial layer, suggesting that close proximity to the *Chlamydia*-infected epithelium may enhance their protective capacity. Such positioning could facilitate the delivery of effector cytokines directly to infected epithelial cells or neighboring immune cells, thereby promoting bacterial clearance. Alternatively, as discussed above, localization near the epithelium may enable CXCR6⁺ CD4 T cells to exert direct cytotoxic activity against infected cells.

Our studies so far have identified both transcription factor BHLHE40 and surface marker CXCR6 as hallmarks of polyfunctional CD4 T cells in *Chlamydia* infection in mice. Several observations support the idea that BHLHE40 may directly regulate the generation of the CXCR6+ effector CD4 T cells. First, *Bhlhe40*-deficient mice exhibited a marked reduction in CXCR6+ cells accompanied by expansion of stem-like CD4 T cells and diminished polyfunctional responses. Second, both *Bhlhe40* deficiency and depletion of CXCR6-expressing cells resulted in impaired bacterial clearance during the late phase of *Chlamydia* infection. Remarkably, a recent study has demonstrated direct binding of BHLHE40 to the regulatory regions within the *Cxcr6* locus ^43^. Together, these findings support a model that TCR/CD28-induced BHLHE40 expression orchestrates an effector CD4 T cell differentiation program that promote polyfunctional CD4 T cell production, in part through induction of CXCR6 expression.

Whether CXCR6+ CD4 T cells persist within the memory CD4 T cell pool following *Chlamydia* clearance remains unknown. In multiple tissues, including the lung, skin, liver, and brain, CXCR6 marks tissue-resident CD8 T cells that contribute to local immune surveillance ^21,33,44,45^. It is therefore tempting to speculate that CXCR6+ CD4 T cells may also be preferentially retained within the FRT following resolution of *Chlamydia* infection. This hypothesis is supported by the observation that CXCR6+ CD4 T cells are enriched in the CD4 MLCs at day 21 post infection (Fig. 5), a time point where these MLC structures begin to form ^31^. Future studies examining the kinetics of CXCR6+ CD4 T cell responses and the pathways governing MLC development will be needed to determine whether CXCR6 expression identifies long-lived tissue resident memory populations. In summary, our study establishes CXCR6 as a marker of protective polyfunctional CD4 T cells and suggest that induction of CXCR6+ effectors may represent a useful correlate of vaccine-induced immunity against *Chlamydia*.

## Acknowledgements

We thank Andrea Harris at the UAMS Flow Cytometry Core for assistant with our flow cytometry experiments. This study was supported by grants from the National Institutes of Health to LXL. (AI139124, AI180670 and GM103625), and American Association of Immunologists Careers in Immunology Fellowship to MABM.

## Author Contributions

MABM and LXL conceived the study, acquired funding and wrote the manuscript. MABM and YK performed the experiments. MABM, QL, and LXL analyzed the data.

## Competing Interests

The authors declare no competing interests.

## Data Availability

The single-cell RNA sequencing dataset analyzed in this study was previously published and is available through the NCBI Gene Expression Omnibus (GEO) under accession number GSE253394.

## Notes

### Competing Interest Statement

The authors have declared no competing interest.

